# A Concept-Driven Disentanglement Framework for Interpretable Graph Neural Networks in Structure-Function Coupling

**DOI:** 10.64898/2025.12.15.694495

**Authors:** Devin Setiawan, Sumaiya Shomaji, Joaquín Goñi, Arian Ashourvan

## Abstract

Graph Neural Networks (GNNs) achieve state-of-the-art performance in predicting brain functional connectivity (FC) from structural connectivity (SC), yet their “black-box” nature limits interpretability and scientific utility. We present a concept-driven disentanglement framework that builds inherently interpretable GNNs for quantitative hypothesis testing of structure-function relationships. The framework employs an ensemble of GNN branches, each architecturally biased to learn from a predefined structural concept (e.g., strong vs. weak connections) by processing a filtered version of the SC graph. This design enforces verifiably disentangled node embeddings, ensuring each branch captures a distinct structural feature. Using SHAP (Shapley additive explanations), we quantify the predictive contribution of each concept and assess its statistical significance against null distributions. Our framework demonstrates high predictive accuracy for FC, achieving a group-level correlation coefficient of 0.91 on a public human connectome dataset, while simultaneously yielding interpretable neuroscientific insights. This interpretable-by-design methodology bridges the gap between predictive power and scientific transparency, enabling deep learning models to provide mechanistic insights into the brain’s network organization.

## 1. Introduction

A central goal in neuroscience is to understand the relationship between the brain’s physical architecture and its dynamic patterns of activity, a principle known as structure-function coupling [1]. Structural connectivity (SC) is derived from diffusion magnetic resonance imaging (dMRI), which characterizes tissue microstructure and structural pathways [2]. It provides the anatomical scaffold that shapes and constrains neural activity into canonical functional networks [3], [4]. These functional connectivity (FC) networks, typically measured as correlated activity in functional MRI signals, in turn support cognitive and perceptual functions [5], [6]. However, the mapping is not a simple one-to-one correspondence, but a complex, non-linear relationship arising from high-order, polysynaptic interactions across the structural network [7].

Understanding this coupling is of critical clinical and scientific importance, as atypical patterns are a known hallmark of numerous neurological and psychiatric disorders, and variations are linked to cognitive performance and brain development [8], [9], [10], [11], [12], [13].

Numerous computational approaches have been developed to model the mapping from SC to FC, revealing a trade-off between mechanistic interpretability and predictive performance. Network and biophysical models represent an early class of highly interpretable methods. They predict FC by simulating neural dynamics across the structural connectome, using either simple communication rules or systems of differential equations [14], [15], [16], [17]. While these models provide valuable mechanistic insight, their predictive accuracy is often modest. Similarly, spectral graph models leverage the eigenmodes of the structural graph Laplacian to predict FC, providing an elegant yet abstract connection between the structure’s resonant modes and its function [18]. In pursuit of higher accuracy, the field has increasingly adopted complex, data-driven machine learning models. Multilayer Perceptrons were shown to significantly improve predictive performance by learning a non-linear mapping from SC to FC, establishing a new benchmark for accuracy [19], [20]. More recently, Graph Neural Networks (GNNs), which are architecturally suited to learn from the connectome’s topology, have further improved performance, achieving state-of-the-art results [21], [22]. However, this progression towards higher accuracy has come at the cost of transparency. Their application in SC-FC coupling often fails to evaluate how topological information contributes to the final predictions. These high-performing deep learning models inherently operate as “black boxes,” learning complex, entangled representations in their latent spaces. Consequently, a critical gap emerged where the most powerful predictive models for SC-FC coupling are the least equipped to provide clear, human-understandable insights or to facilitate the testing of specific neuroscientific hypotheses.

The challenge of model opacity has given rise to a significant body of research in explainable AI for GNNs (XAI) [23]. The predominant approaches are post-hoc, aiming to explain a model’s decisions after it has been trained. These methods, such as GNNExplainer, PGExplainer, and SubgraphX, have proven powerful for identifying salient input features (e.g., a minimal set of critical edges, influential subgraph, etc) that contribute to a specific prediction [24], [25], [26]. However, a fundamental limitation of post-hoc analysis in this context is that it attempts to rationalize a model that has already learned from a complex mixture of information, resulting in entangled latent representations. While these methods can highlight which parts of the input graph are statistically significant, the resulting explanations do not inherently map to predefined, human-understandable neuroscientific concepts. They also do not fully explain why a particular subgraph is important from a domain perspective. This leaves a critical research gap for a methodology that can bridge the model’s internal logic with explicit, hypothesis-driven scientific inquiry.

To address this gap, we move beyond post-hoc rationalization to propose an inherently interpretable framework. Our goal is to allow Graph Neural Network architectures, which have been proven to perform well for this task, to be modified to explicitly encode predefined structural concepts, thereby enabling a verifiable and quantitative framework for interpretable structure-function coupling. We achieve this through a concept-driven disentanglement framework that utilizes an ensemble of specialized GNNs, where each branch is architecturally biased via input graph filtering to process a distinct, user-defined structural concept. We demonstrate that this approach produces verifiably disentangled node embeddings while maintaining predictive performance comparable to state-of-the-art models. Furthermore, we present a complete workflow for using this framework as a tool for quantitative hypothesis testing. By applying SHAP analysis and comparing against null distributions from random concepts, we show that our model learns non-trivial, neuroscientifically plausible relationships, such as prioritizing the less prevalent but more informative strong connections over weak ones [27]. This “interpretable-by-design” methodology provides a tool for quantifying the predictive importance of specific structural properties, allowing for more insightful applications of deep learning in neuroscience.

## 2. Methodology

This section outlines the methodological framework developed to implement and evaluate our concept-driven, interpretable GNN for structure–function coupling. We describe the dataset and preprocessing steps, the formulation of the GNN model, and the architectural design of the proposed disentanglement framework. Subsequent subsections detail the training setup, workflow for quantitative hypothesis testing, and procedures for verifying conceptual disentanglement.

### A. Dataset and Preprocessing

The data used in this study were obtained from the MICA-MICs (Microstructure-Informed Connectomics) dataset, an open-access resource for connectomics research [28]. We utilized a cohort of 50 healthy adult subjects for which both structural and functional connectivity data were available. The connectomes were parcellated using the Schaefer 100-region atlas, resulting in connectivity matrices of size 116×116 (100 cortical regions and 16 subcortical regions) for each subject. All data were acquired and processed using the micapipe pipeline [29].

The raw SC and FC matrices provided in the dataset were supplied as upper triangular matrices. For each subject, we first constructed the corresponding full, symmetric connectivity matrix. To ensure numerical stability during model training and to standardize the input feature range across subjects, we applied a subject-wise min-max normalization to both the SC and FC matrices. For a given subject’s matrix *M* (either SC or FC), the normalized matrix *M*_*norm*_was calculated as:

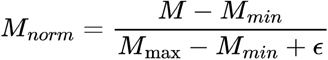

where *M*_*min*_ and *M*_*max*_ are the minimum and maximum values within that subject’s matrix, and *ϵ* is a small constant (10^−8^) to prevent division by zero. This procedure scales all connection weights to the range [0, 1] for each subject independently.

### B. GNN Formulation for Structure-Function Coupling

Following recent state-of-the-art approaches by Chen et al., we model the brain connectome as a graph, *G =* (*V, E*), where the set of nodes *V* represents *N* brain regions of interest (ROIs), and the set of edges *E* represents the structural connections between them. For a given subject, the SC matrix *S* ∈ *R*^*N*×*N*^ provides the edge weights for this graph. Initial node features are represented by an identity matrix *X* ∈ *R*^*N*×*N*^.

The backbone of the predictive model is a Graph Convolutional Network (GCN), a type of GNN that learns node embeddings by aggregating and transforming features from neighboring nodes [30]. The layer-wise propagation rule for a GCN is as follow:

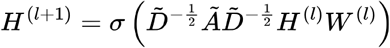

where *Ã = A +I* is the adjacency matrix with added self-loops, 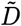 is its degree matrix, *H*^(*l*)^ are the node activations at layer *l* with initial state *H*^(0)^ =*X, W*^(*l*)^ is a trainable weight matrix, and *σ* is an activation function. After *L* layers of convolution, the GCN produces a final node embedding matrix *E* ∈ *R*^*N*×*d*^, where *d* is the embedding dimension. The functional connectivity (FC) value between any two nodes, *u* and *v*, is then predicted by a Multi-Layer Perceptron (MLP) that takes the concatenated embeddings of the node pair, [*E*_*u*,_ *E*_*v*_], as input [31].

### C. Concept-Driven Disentanglement Framework

To overcome the inherent lack of interpretability in standard GNNs, which learn entangled representations, we introduce a framework for achieving explicit conceptual disentanglement by architectural design. The overall architecture is shown in Fig. 1.

**Figure 1.**
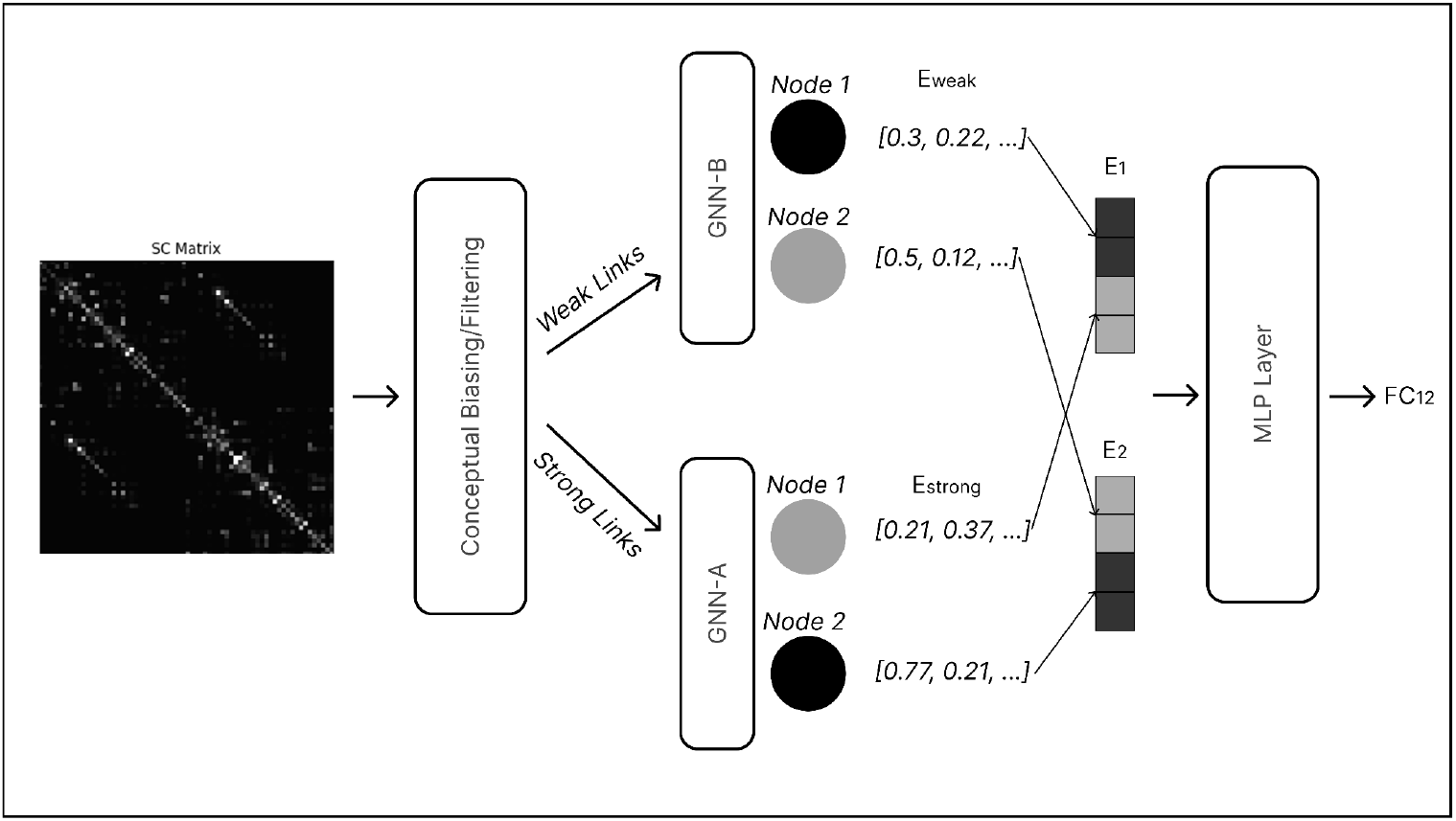
Overall Architecture. The overall architecture of the proposed model. An input Structural Connectivity (SC) matrix undergoes “Conceptual Biasing,” where it is filtered into multiple, conceptually distinct graph views (e.g., “Weak Links,” “Strong Links”). Each view is processed by a separate, specialized GNN branch (GNN-A, GNN-B) to produce a concept-specific node embedding. These specialized embeddings are concatenated into a final, disentangled representation for each node. Two disentangled embeddings are concatenated and processed by a terminal MLP layer to produce an FC prediction.

The framework is predicated on “conceptual biasing,” a method of filtering the input SC matrix based on predefined, human-understandable structural properties. We formally define a structural “concept” as a transformation function, *T*_*k*_ (·), that generates a biased SC matrix *S*_*k*_ = *T*_*k*_ (*S*), where *S*_*k*_ contains a specific subset of edges from the original matrix *S*. We list all the formal definitions for the conceptual biasing transformations (*T*_*k*_) used in the experiments in Table 1. Each concept partitions the set of existing positive-weighted connections, *E =* {(*i, j*) | *S*_*ij*_ > 0 }, into two mutually exclusive sets. Concepts differ in whether their thresholds are defined at the subject level (individual variability) or group level (population distribution). This is explicitly shown in the mathematical notation below.

**Table 1.**
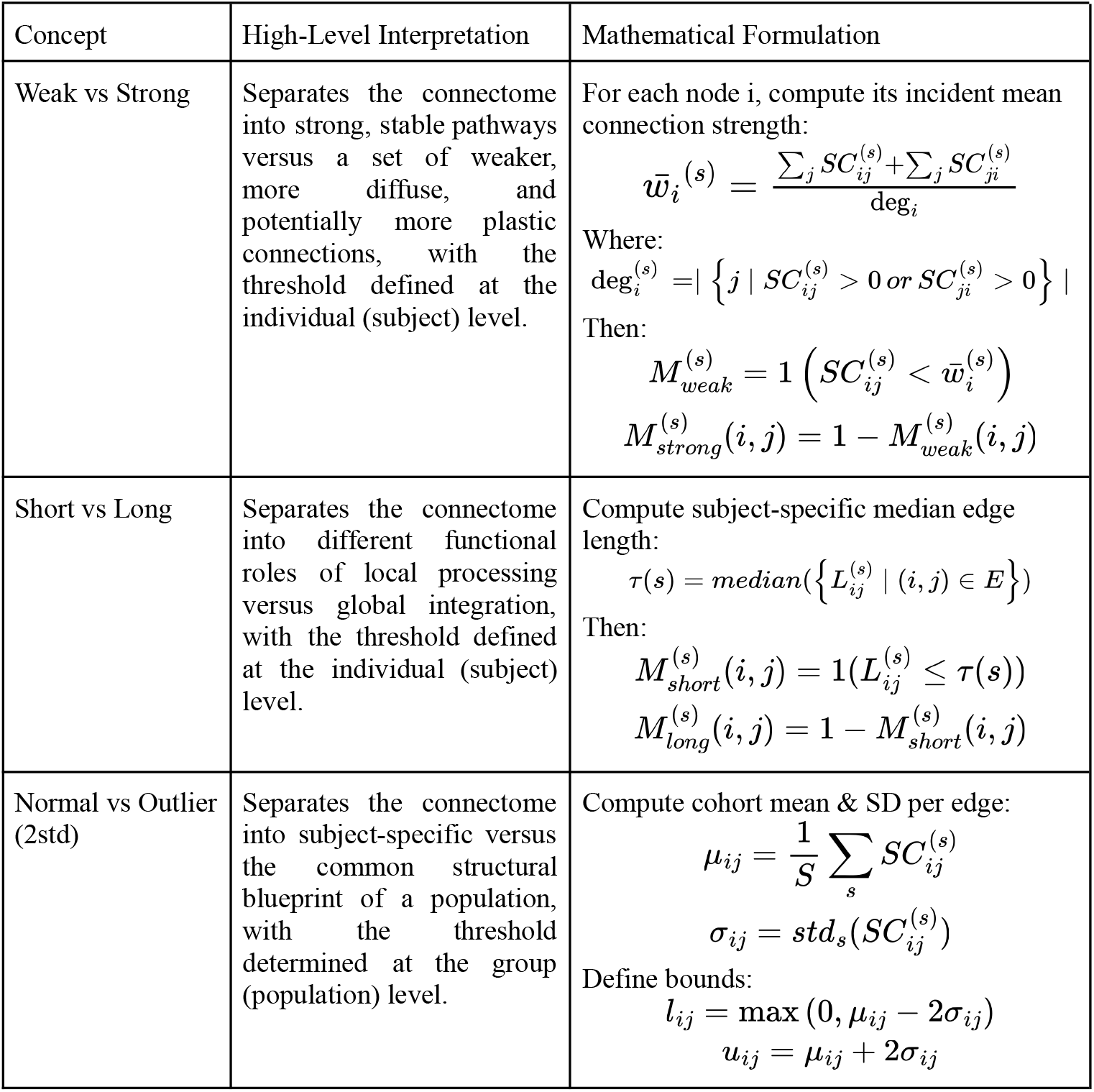
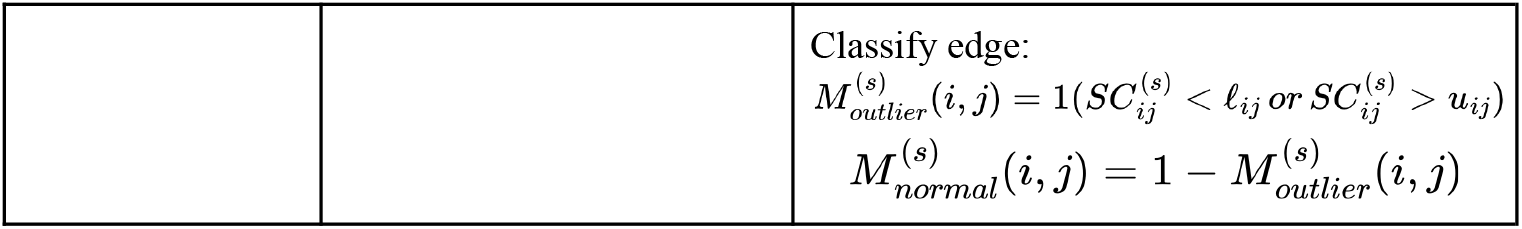
List of Concepts.

Let

1. *SC*^(*s*)^ ∈ *R*^*N*×*N*^ : structural connectivity matrix for subject *s*
2. *L*^(*s*)^ ∈ *R*^*N*×*N*^ : connection-length matrix for subject *s*
3. *E =* {(*i, j*) | *S*_*ij*_ > 0 } : set of existing edges
4. *μ ij, σ ij* : population-level mean and standard deviation across subjects

The binary mask for concept *c* and subject *s* is:

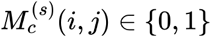

The core of the framework is an ensemble of specialized GNN branches. Instead of a single GNN processing the full SC matrix, each branch is fed one of the conceptually biased matrices *S*_*k*_. This forces each branch to learn a node embedding, *E*_*k*_, that is representative of its specific conceptual input. These specialized embeddings are then concatenated to form a final, disentangled node embedding for each brain region before being passed to the prediction MLP.

### D. Detailed Ensemble Architecture

The specific implementation of our 2-feature ensemble is detailed in Fig. 2. Each SingleGNN branch consists of a two-layer GCN. The first layer maps the initial 116-dimensional node features to a 32-dimensional hidden space. The second layer further processes these features, outputting a final 32-dimensional concept-specific embedding. For a given node, the 32D short-connection embedding and the 32D long-connection embedding are concatenated to produce a 64D disentangled node representation. To predict the FC between a source node u and a target node v, their respective 64D embeddings are concatenated, resulting in a 128D vector. This vector is then processed by a two-layer MLP (with a 128-dimensional hidden layer) to regress the final scalar FC value. ReLU activations are used throughout the network, and the model was trained using the Adam optimizer. The model was trained on the dataset using a 70/30 train-test split. We used the Adam optimizer with an initial learning rate of 0.01 and the Mean Squared Error (MSE) loss function with ‘mean’ reduction. A ReduceLROnPlateau learning rate scheduler was applied, reducing the learning rate by a factor of 0.1 after 5 epochs of no validation loss improvement. Data was loaded with a training batch size of 2 (shuffled) and a testing batch size of 1 (unshuffled), and the model was trained for 200 epochs.

**Figure 2.**
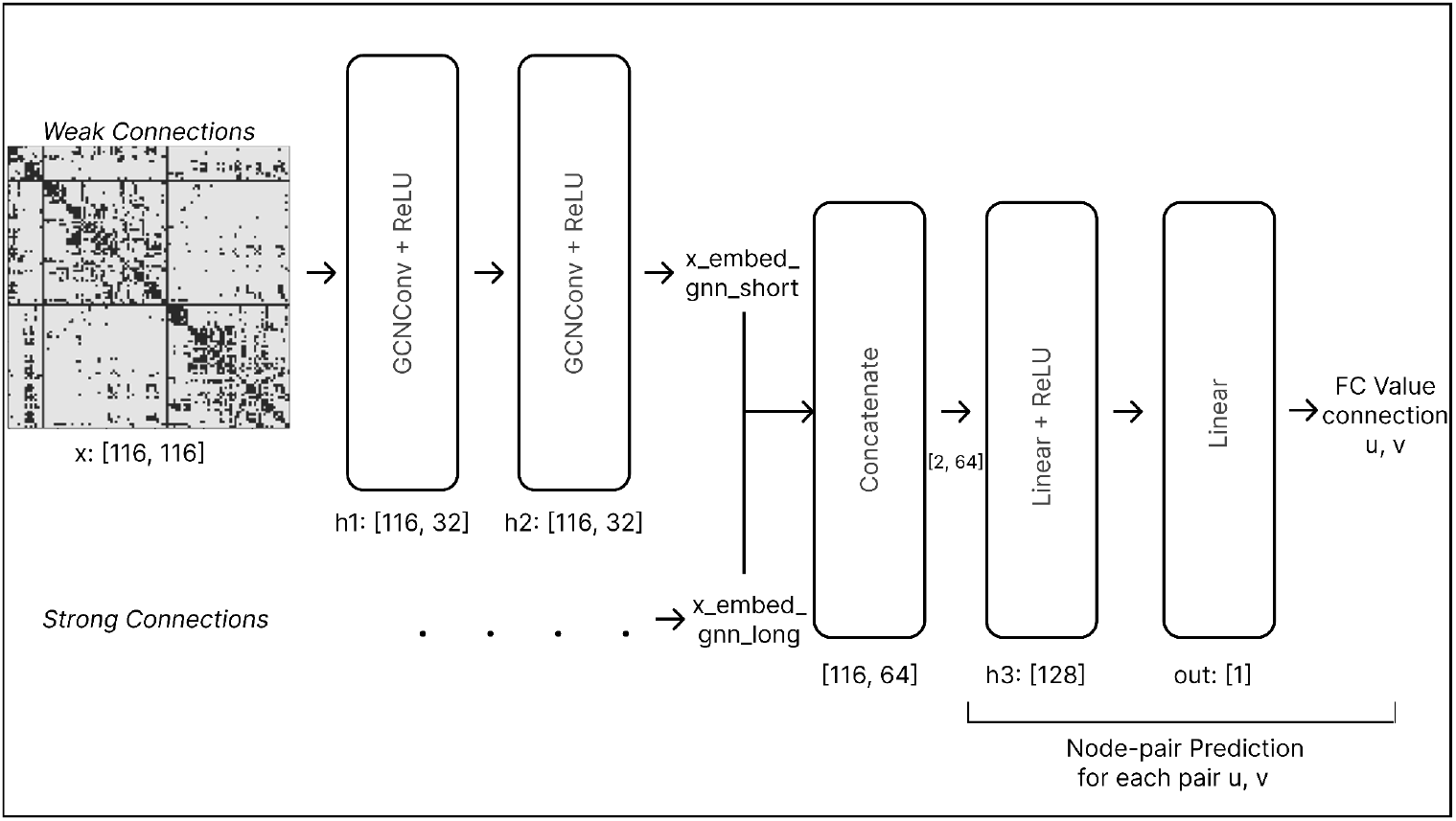
Disentangle GNN Ensemble Architecture. A schematic detailing the flow of tensor dimensions through a two-branch ensemble. A conceptually biased SC matrix (e.g., “Weak Connections”) and initial node features (x: 116×116) are input to a branch. This branch consists of a two-layer GCN, which produces a concept-specific 32-dimensional embedding (h2: 116× 32). The embeddings from the “short” and “long” branches are concatenated for each node, resulting in a 64D disentangled representation (116×64). For predicting the FC value between a node pair u, v, their 64D embeddings are concatenated into a 128D vector (2×64) and regressed to a single scalar output by a two-layer MLP.

A key challenge of conceptual biasing is that partitioning the SC matrix can result in a significant imbalance in the number of edges representing each concept (e.g., a 95/5 split for “Normal” vs. “Outlier” connections). To prevent the model from preferentially learning from the concept that has more data and to ensure a fair comparison in the subsequent importance analysis, we implemented a weighted Mean Squared Error (MSE) loss function. The weight for each edge’s loss was determined dynamically within each training batch based on the inverse frequency of its corresponding concept. For a batch containing *N*_*batch*_ edges, let *N*_*c*1_ and *N*_*c*2_ be the number of edges belonging to Concept 1 and Concept 2, respectively. The weights for each concept are calculated as:

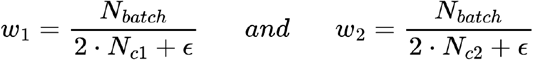

where *ϵ* is a small number (10 ^−6^) for numerical stability. A weight vector *W*_*loss*_ is then constructed, assigning the appropriate weight (*w*_1_ *or w*_2_) to each edge in the batch based on its conceptual class. The final weighted loss L for the batch is the mean of the element-wise product of the per-edge MSE and their assigned concept weights:

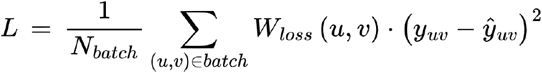

### E. Workflow for Quantitative Hypothesis Testing

The proposed framework enables a systematic workflow for testing neuroscientific hypotheses, as illustrated in Fig. 3. The process begins with defining a hypothesis as a comparison between two or more structural concepts (e.g., “Are strong connections more predictive of FC than weak connections?”). An ensemble GNN is then trained with branches architecturally biased to these mutually exclusive concepts. After training, the predictive importance of each disentangled embedding component is quantified. We employed SHAP (Shapley additive explanations), a game-theoretic approach to explain the output of any machine learning model. We used the GradientExplainer variant, an extension of Integrated Gradients to calculate the contribution of each dimension of the concatenated node embeddings for every predicted edge in the test set [32].

**Figure 3.**
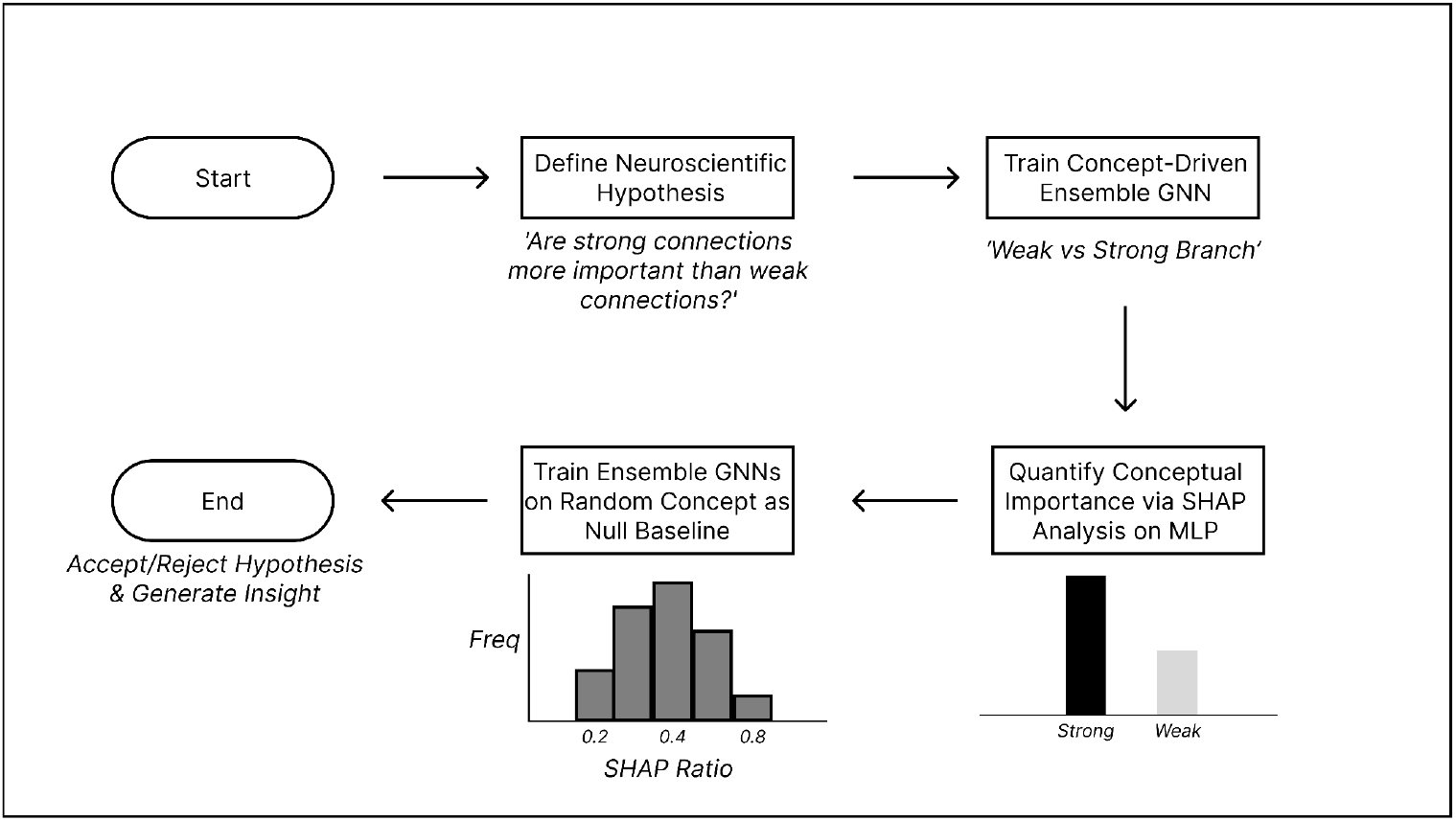
Workflow for Quantitative Hypothesis Testing. (1) A hypothesis is defined (e.g., comparing strong vs. weak connections). (2) A concept-driven ensemble GNN is trained with branches architecturally biased to these concepts. (3) The predictive importance of disentangled embedding components is quantified using SHAP analysis. (4) The resulting importance ratio is compared against a null distribution generated by training identical ensembles on random concepts with the same data prevalence. (5) This comparison allows for the acceptance or rejection of the initial hypothesis.

To ensure that the quantified importance of each concept was not an artifact of its prevalence in the dataset, we compared the observed SHAP importance ratio between concept branches to a null distribution. This null distribution was obtained through a permutation-based procedure with 100 iterations, in which the same ensemble architecture was retrained 100 times on randomized concept assignments that preserved the original data prevalence (e.g., maintaining an 81/19 split of edges). For each iteration, SHAP importance ratios were recalculated to form the null distribution. We then computed a one-tailed p-value for each concept’s observed SHAP ratio. This p-value represents the proportion of the 100 ratios in the null distribution that were as extreme or more extreme than the observed ratio (in the hypothesized direction). A concept’s importance was deemed statistically significant if its observed SHAP ratio lay outside the 95th percentile of this null distribution (*p* < 0.05). This permutation testing procedure formally enabled acceptance or rejection of the initial hypothesis.

### F. Verification of Conceptual Disentanglement

To empirically validate that our ensemble architecture produces disentangled node embeddings, we performed a targeted perturbation analysis on a trained ‘Short vs Long’ model. The analysis was designed to test whether each GNN branch was selectively sensitive only to its designated conceptual input. The procedure involved perturbing the input test data by adding noise to the input labeled to a particular concept branch. Following each perturbation, we passed the modified test data through the trained model to generate new node embeddings. We quantified the change by computing the Euclidean distance between the original embeddings and the perturbed embedding for each branch, averaged across all nodes and subjects. An Euclidean distance of zero would indicate perfect stability (no change).

## 3. Results

This section presents the empirical results of our study. We first establish the predictive performance and topological validity of our models. We then demonstrate the framework’s utility for quantitative hypothesis testing by analyzing the predictive importance of several predefined structural concepts.

### A. Model Performance and Topological Dependence

To validate our implemented models, we first benchmarked their predictive performance against established methods from the literature. As shown in Fig. 4(a), our Single GNN Baseline achieved a group-level Pearson correlation of r=0.95 and an average individual-level correlation of r=0.81. The 2-feature Ensemble GNN, designed for interpretability, achieved a group-level r=0.91 and an individual-level r=0.78. These results were comparable to the state-of-the-art GNN model (r=0.95 group, r = 0.69 individual), confirming the high predictive accuracy of our architectures.

**Figure 4.**
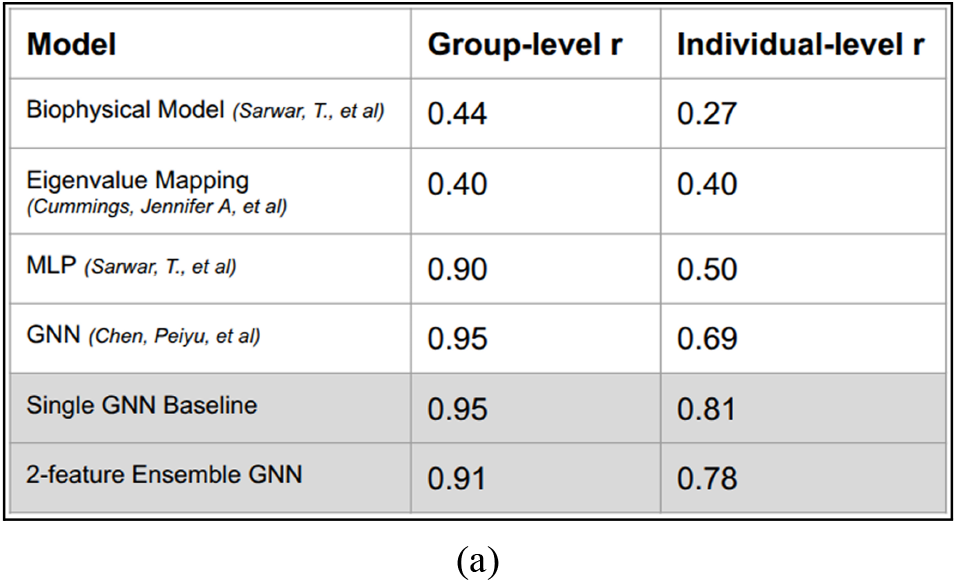
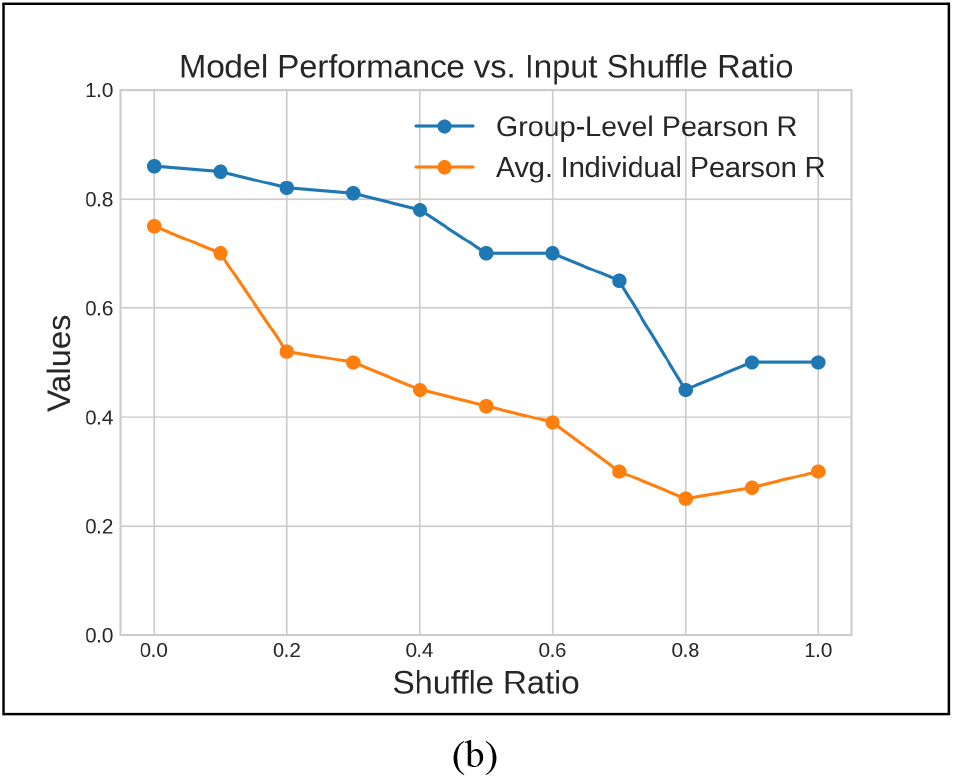
Model Performance and Topological Dependence. (a) Predictive performance of our Single GNN Baseline and 2-feature Ensemble GNN compared to benchmark models from the literature. Our models achieve state-of-the-art accuracy and a favorable accuracy trade-off for the interpretable ensemble GNN. (b) A null model experiment validating that the model learns from the specific topology of the SC. Performance degrades as an increasing fraction of the SC edge weights are randomly shuffled, confirming that the model is not learning from simple statistical properties of the input but from its graph structure.

To confirm that the models learned from the underlying graph structure, a null model experiment was conducted by progressively shuffling the input SC edge weights. Fig. 4(b) shows a degradation in both group-level and individual-level Pearson correlation as the ratio of shuffled edges increased. At a shuffle ratio of 0.8, the average individual performance dropped to r=0.27, demonstrating that the models’ predictive power is dependent on the topology of the structural connectome.

### B. Conceptual Disentanglement Verification Result

A core claim of our framework is that the conceptual biasing forces the GNN branches to learn disentangled representations. We tested this empirically using the targeted perturbation analysis detailed in the Methodology section. The results, summarized in Table 2, confirm a highly successful disentanglement. Perturbing one conceptual input caused a disruption in its corresponding embedding, while the embedding from the unperturbed branch remained entirely unaffected. This verification provides a solid foundation for attributing the model’s predictive importance to specific, isolated structural concepts.

**Table 2.** Perturbation Analysis Result.

| Perturbation Target | Avg. Euclidean Distance of $E_{short}$ | Avg. Euclidean Distance of $E_{long}$ |
| --- | --- | --- |
| Short Connection | 0.2438 | 0.0000 |
| Long Connection | 0.0000 | 0.0320 |

### C. Quantifying the Predictive Importance of Weak vs. Strong Connections

We first applied our framework to test the hypothesis regarding the relative importance of weak versus strong structural connections. The SC matrix was partitioned into two conceptual inputs based on connection strength, with approximately 81% of positive-weighted edges classified as ‘weak’ and 19% as ‘strong’.

The predictive importance of each concept branch was quantified using SHAP analysis on the trained ensemble. Fig. 5 shows the null distribution comparison between random splitting of the input data and a neuroscientifically meaningful concept (Weak vs Strong). The strong connections, despite representing a minority of the input data (19%), accounted for the majority of the predictive importance, with a sum of mean absolute SHAP values of approximately 0.27 compared to less than 0.01 for the weak connections. To assess the statistical significance of this finding, the observed SHAP contribution ratio for the neuroscientifically defined concept (75.5) was compared against the null distribution from the permutation test (N=100 iterations). The observed ratio was found to be a significant outlier. It was more extreme than all 100 ratios generated for the null distribution (null distribution: Mean=0.42, SD=0.27), corresponding to a p-value of p < 0.001. This is well below our pre-defined significance.

**Figure 5.**
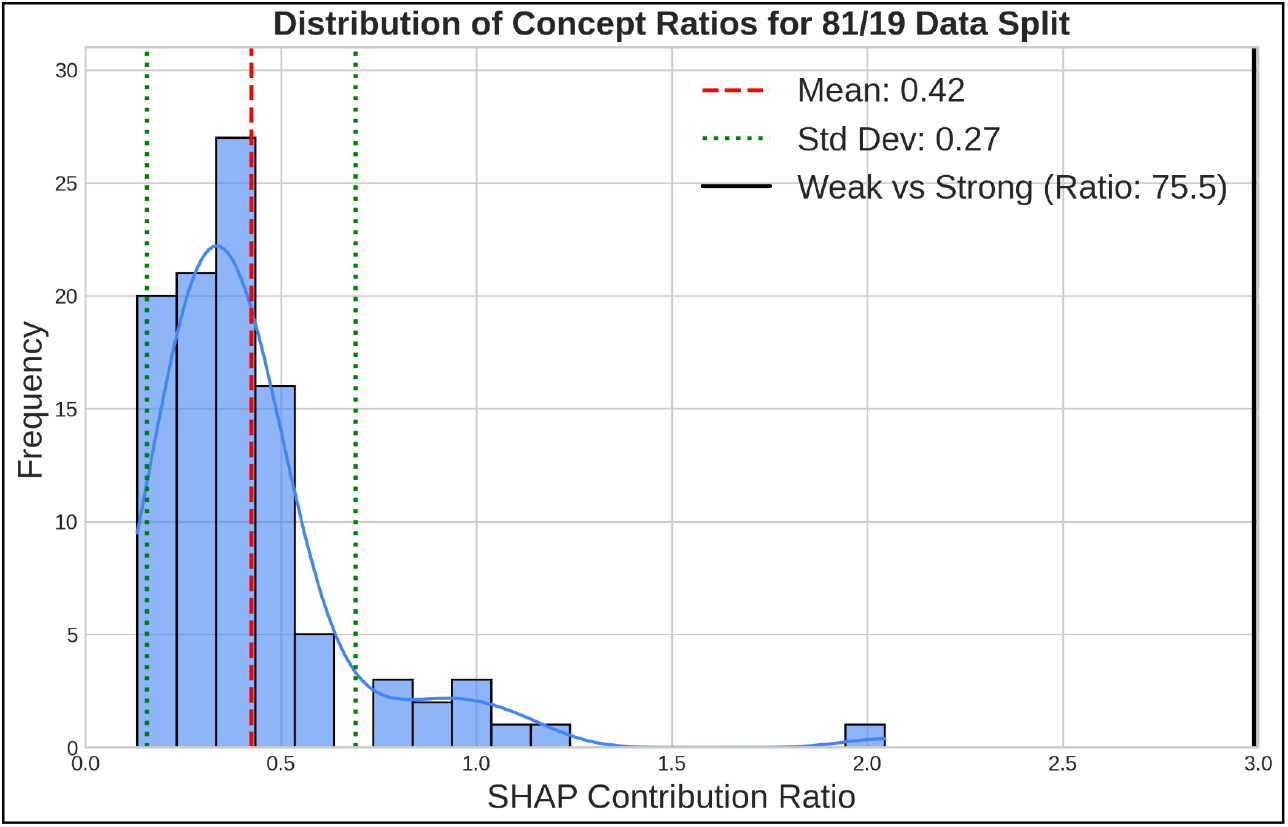
Quantifying Predictive Importance of Weak vs Strong Connections. Strong connections represent a minority of the input data (∼19%), while weak connections represent a majority of the input data (∼81%). The histogram is a null distribution of SHAP contribution ratios, generated by training ensembles on random 81/19 splits of the edge data. The observed SHAP ratio for the “Weak vs. Strong” concept (Ratio: 75.5) is a clear outlier. Although strong connections are only made up of a minority of the input data, they account for most of the predictive outcome of the model. The model has learned a meaningful relationship that is statistically significant beyond what would be expected by a random data partition.

### D. Framework Generalizability for Other Structural Concepts

To demonstrate the flexibility of the framework for testing diverse hypotheses, we evaluated two additional pairs of structural concepts.

First, we examined the importance of “Short” versus “Long” connections, defined by a median split of fiber lengths. The SHAP analysis, shown in Fig. 6(a), revealed that the model attributed a disproportionately large predictive importance to short-range connections. The observed SHAP ratio of 0.17 for this concept was compared against its null distribution from the permutation test (N=100 iterations). The observed ratio was found to be a significant outlier. It fell into the lowest 5% of the 100 ratios generated for the null distribution (null distribution: Mean = 1.13, SD = 0.56), corresponding to a one-tailed p-value of p < 0.05. This meets our pre-defined significance level.

**Figure 6.**
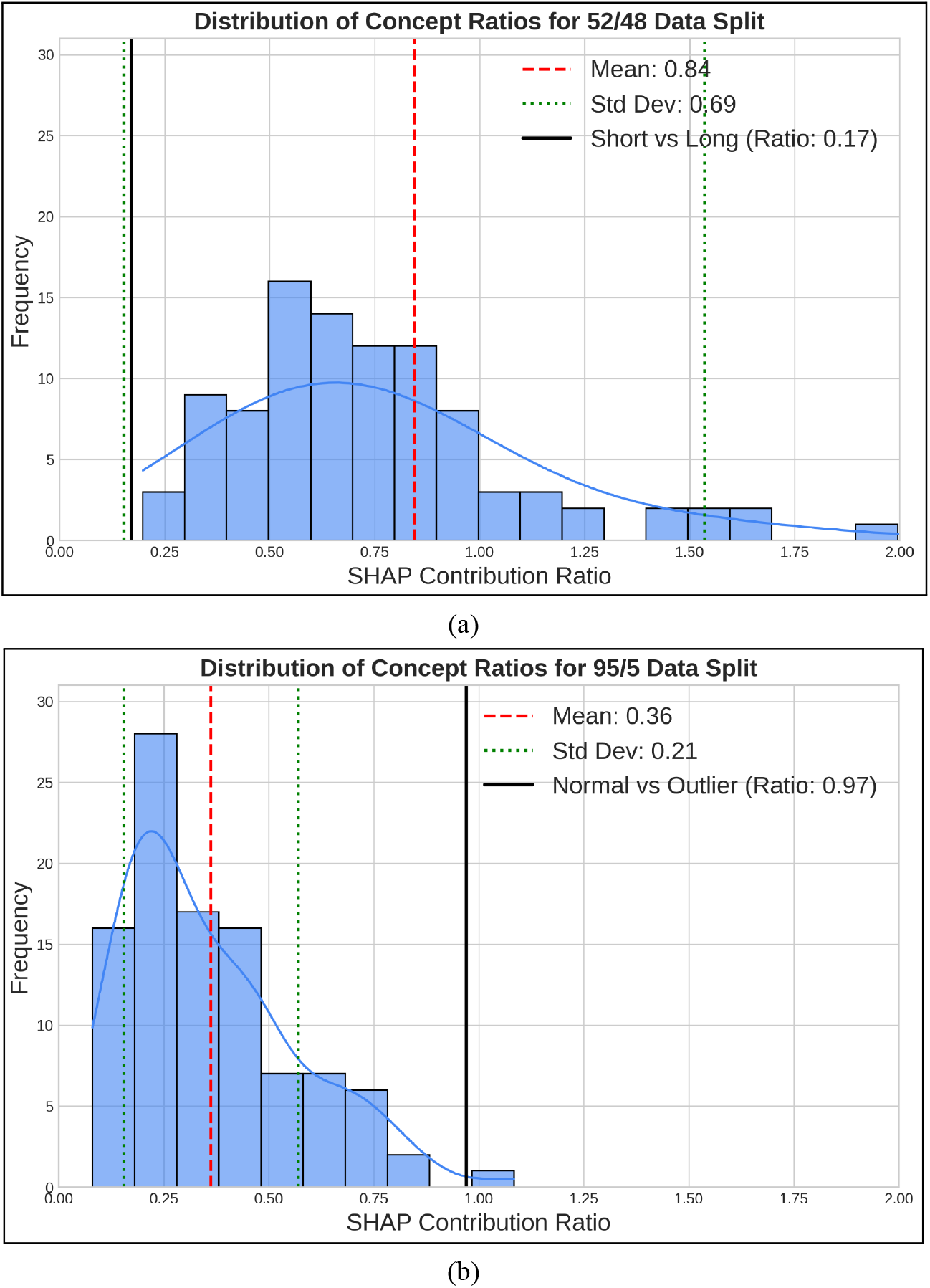
Assessing the Importance of Other Structural Concepts. (a) We biased two GNN branches using the median split of connection lengths. The model attributes a disproportionately large predictive importance to short-range connections. This is evidenced by an observed SHAP ratio (Ratio: 0.17) that is a clear outlier from its random baseline expectation. (b) We defined “outlier” connections as those with weights greater than two standard deviations above the mean (∼5% of edges). The model learns to assign a surprisingly balanced importance. It attributes a much larger predictive role to the rare “outlier” connections than would be expected by chance, as shown by its SHAP ratio (Ratio: 0.97) being a significant outlier above its random baseline.

Second, we evaluated “Normal” versus “Outlier” connections, where outliers were defined as connections with weights greater than two standard deviations above the mean, constituting approximately 4% of the data. The results in Fig. 6(b) show that the model assigned a surprisingly balanced importance to these concepts. The observed SHAP ratio of 0.97 for this concept was compared against its null distribution from the permutation test (N=100 iterations) for the 95/5 split. The observed ratio was found to be a significant outlier. It exceeded the 95th percentile of the 100 ratios generated for the null distribution (null distribution: Mean = 0.36, SD = 0.21), corresponding to a one-tailed p-value of p < 0.05. This meets our pre-defined significance threshold, indicating that the rare outlier connections contributed disproportionately to the prediction.

## 4. Discussion

This section interprets the implications of our findings, evaluating the utility of our interpretable-by-design GNN for modeling structure–function coupling. We begin by summarizing the principal findings, focusing on the model’s ability to achieve state-of-the-art predictive accuracy while maintaining verifiably disentangled representations. We then contrast this methodological approach with standard post-hoc explainability. Subsequent subsections examine the specific neuroscientific insights revealed by the model, and conclude with an assessment of methodological limitations and directions for future research.

### A. Principal Findings

Our findings demonstrate that an interpretable-by-design GNN can achieve state-of-the-art performance in predicting FC from SC while maintaining conceptual transparency. Unlike traditional “black-box” GNNs that conflate multiple structural factors into a single latent representation, our architecture successfully disentangled predefined structural concepts such as strong versus weak, short versus long, and normal versus outlier connections into independent embedding spaces. The empirical verification via perturbation analysis confirmed that each branch of the ensemble was selectively sensitive to its designated conceptual input, validating the intended disentanglement.

Crucially, this interpretability did not come at a cost to predictive accuracy. The ensemble achieved an individual-level correlation (r = 0.78) comparable to the best-performing entangled GNNs (r = 0.81), demonstrating that meaningful interpretability and high performance can coexist. The model also passed a stringent topology-dependence test, in which random shuffling of structural edges progressively degraded prediction accuracy. This outcome supports the notion that the model genuinely leverages connectomic topology rather than superficial statistical regularities, echoing earlier findings that FC is deeply constrained by the brain’s structural architecture [1], [14], [16].

### B. Interpretable-by-Design vs. Post-Hoc Explainability

This work’s “interpretable-by-design” methodology not only offers a direct response to the limitations of post-hoc explainability, but also a methodological bridge between deep learning and theory-driven computational modeling. Our framework reframes the GNN architecture as an explicit embodiment of competing neuroscientific theories. This approach contrasts sharply with post-hoc explainability, which attempts to rationalize a model after it has learned an entangled latent representation [23], [24], [25], [26]. Post-hoc graph explanation methods are designed to identify the importance of individual features (e.g., specific edges or nodes). This is fundamentally different from our goal of quantitative hypothesis testing. In our framework, interpretability is embedded directly into the architecture. Each GNN branch is architecturally constrained from the start to learn from separate, human-understandable concepts. Our perturbation analysis empirically confirmed that this design creates verifiably disentangled subspaces. This validation confirms the conceptual integrity of our disentangled subspaces, a guarantee that is not (and cannot be) offered by post-hoc methods operating on entangled models. SHAP is therefore not used to explain a black box, but to quantify the predictive contribution of isolated, concept-specific embeddings [27]. This enables concept-level explainability: granular enough to test neuroscientific hypotheses (e.g., “strong connections dominate prediction”), yet abstract enough to avoid interpreting meaningless individual embedding dimensions. Ultimately, this framework allows researchers to treat the model itself as a testbed for theory, where architectural choices directly encode theoretical commitments. This is a task for which post-hoc feature-attribution methods are not designed for.

### C. Neuroscientific Implications

The framework’s disentangled design allows direct interpretation of how structural properties shape functional coupling. Three consistent patterns emerged: (i) dominance of strong connections, (ii) disproportionate importance of short-range links, and (iii) substantial predictive value of rare outlier edges.

#### 1. Strong Structural Connections Form the Stable Backbone of Static FC

A key finding of our framework was the disproportionate predictive importance of strong, high-weight structural connections. This result provides insight into the nature of structure-function coupling when modeled with graph neural networks. First, our GNN architecture is inherently a polysynaptic model. Mathematically, our 2-layer GCN architecture is equivalent to aggregating information from two-hop topological neighborhoods, a depth shown in prior optimization studies to maximize predictive accuracy in brain networks [21]. This design allows the model to simultaneously integrate direct (1-hop) and indirect (2-hop) signaling effects [16], [33], [34]. Our model learned that the most informative pathways for predicting functional connectivity are those routed along the “backbone” of strong structural connections. This indicates that regardless of whether the communication is direct or polysynaptic, its predictive contribution is contingent on the integrity of the strong, high-weight tracts that scaffold these paths. This finding supports a specific mechanistic hypothesis about the nature of static FC. By definition, static FC is a time-averaged correlation that aggregates neural activity over the entire scan duration. This averaging process acts as a temporal filter that prioritizes stable, persistent interactions. Here, the distinction between strong and weak connections becomes critical. Co-activations driven by weaker connections are intermittent and transient. When averaged over the full scan duration, these sparse signals are effectively diluted [35], [36]. In contrast, the strong, high-capacity anatomical tracts are thought to support a stable, persistent ‘baseline’ of neural communication [37]. This consistent, underlying covariance does not average out and thus disproportionately defines the final static FC matrix. Therefore, our model learned to isolate the stable structural ‘backbone’ that best predicts this stable, time-averaged functional outcome. Prior SC–FC coupling studies have similarly found that strong anatomical tracts disproportionately contribute to static functional connectivity patterns, supporting the notion that static FC reflects the stable core of high-capacity pathways [1], [14], [19]. Models designed to explain dynamic, moment-to-moment communication may, in contrast, find a greater role for weaker, more flexible pathways, but explaining this dynamic state was not the goal of the current study.

#### 2. Short-Range Dominance is Consistent with Hierarchical Cortical Organization

The model’s preferential weighting of short-range connections is a finding that likely reflects a combination of a core biological principle and a persistent methodological confound. Biologically, it is consistent with hierarchical cortical organization, where dense local circuitry sustains modular specialization and strong intra-community coupling [38], [39]. The predominance of short-range links in explaining static FC thus aligns with the idea that stable functional architecture is rooted in these local, intra-module circuits. This biological interpretation, however, must be acknowledged alongside a well-known confound: spatial autocorrelation in fMRI signals, whereby smoothing and shared physiological noise inflate correlations between neighboring regions [40]. The central challenge is that our model may be learning to use short-range SC connections as a simple, effective proxy to predict distance-confounded FC values. Our null-model tests (Fig. 4b) confirmed that the model is sensitive to SC topology, but this experiment does not and cannot disentangle true biological coupling from this confound. We must therefore be explicit that we have not controlled for this bias. A definitive interpretation would require further analysis, such as quantifying the distance-FC correlation in our data [41], [42]. Therefore, while the model’s finding is consistent with the principle of local organization, this interpretation is limited, and we cannot rule out that this “short-range dominance” largely reflects a learned association with this known data artifact.

#### 3. Outlier Structural Feature Provides Basis for Capturing Individual Variability

Another key finding was the disproportionate predictive importance of the “outlier” branch. These connections, representing a small fraction of the data (approx. 4-5%) where an individual’s structure deviates significantly from the population mean, were found to be highly predictive and statistically significant. This result indicates that accurate prediction of an individual’s FC requires information beyond the common structural blueprint, and that rare, ‘outlier’ connections are disproportionately informative for that prediction. This finding generates a compelling, testable hypothesis that these predictive outlier connections may represent the structural correlates of functional individuality, or ‘fingerprinting’ [43], [44]. We speculate that while ‘normal’ connections form a shared population blueprint, these sparse outliers could represent the idiosyncratic variations that differentiate individuals. We must be clear that our current work does not test this hypothesis. We show that these outlier connections are predictive on average, but we did not test whether these outlier patterns are themselves idiosyncratic (i.e., unique to each individual). Furthermore, we did not correlate individual-level structural outlier patterns with functional individuality metrics. Therefore, we propose this as a key direction for future research. Investigating whether these model-identified predictive outliers are, in fact, the stable, individual-specific structural variations that drive functional uniqueness is a non-trivial but exciting next step.

### D. Limitations and Future Directions

Despite its interpretive strengths, several limitations warrant further exploration. First, the current implementation relies on hard conceptual partitions (e.g., strong vs. weak) defined by thresholding structural properties. This binary separation simplifies interpretability but may obscure the graded or overlapping nature of real connectomic features. Because each GNN branch is restricted to a filtered subset of the connectome, information sharing between concepts is limited. This, in turn, constrained the model’s ability to capture synergistic or emergent relationships among overlapping structural features, such as the co-occurrence of strong and long-range pathways. Future versions should adopt soft conceptual biasing or attention-based weighting to model interactions among concepts, capturing the continuum of structural organization underlying polysynaptic communication [7], [15].

Second, the ensemble’s modular structure increases computational overhead relative to single-branch architectures, highlighting an inherent trade-off between explanatory granularity and model efficiency. Nonetheless, these trade-offs are justified by the framework’s ability to yield causal and mechanistic insight.

Third, the framework’s interpretability depends on a priori concept definitions grounded in current literature. While this facilitates hypothesis testing, it risks reinforcing existing assumptions about structure–function coupling. Hybrid architectures combining predefined and data-driven concept discovery could allow the model to learn novel structural motifs while retaining transparency.

Fourth, our analysis was limited to a normative cohort of healthy adults. Extending this framework to clinical or developmental datasets could test whether disease or maturation alters the predictive importance of specific structural concepts [13]. Our architecture is uniquely suited to quantify competing hypotheses about neuropathology. For instance, in development, we could directly test the “network integration” hypothesis by tracking an expected shift in predictive dominance from short-range to long-range connections as the brain matures [45]. Furthermore, in conditions like schizophrenia, the model could quantify a hypothesized disruption in the strong-vs-weak connection balance, testing theories of inefficient network organization [46], [47]. In cases of neurodegeneration like Alzheimer’s disease, we could even test for compensatory mechanisms, hypothesizing that the predictive importance of the “outlier” branch would increase as the brain attempts to reroute function around compromised “normal” pathways [48], [49]. Such applications would move beyond prediction toward mechanistic inference, enabling interpretable GNNs to serve as tools for understanding how structural disruptions reshape functional dynamics.

## 5. Conclusion

We have presented a concept-driven disentanglement framework that successfully bridges the gap between high-performance GNNs and interpretable scientific models for brain structure-function coupling. By conceptually biasing an ensemble of GNNs with predefined structural concepts, we produced verifiably disentangled node embeddings without a significant loss in predictive accuracy. We demonstrated the framework’s utility as a novel tool for quantitative hypothesis testing, revealing the concept-dependent priorities the model learns from the connectome. This “interpretable-by-design” methodology represents a significant step towards making advanced deep learning models more transparent, trustworthy, and ultimately more useful for scientific discovery in neuroscience and beyond.

## References

[1] L. E. Suárez, R. D. Markello, R. F. Betzel, and B. Misic, “Linking Structure and Function in Macroscale Brain Networks,” Trends Cogn. Sci., vol. 24, no. 4, pp. 302–315, Apr. 2020, doi: 10.1016/j.tics.2020.01.008.

[2] H. Johansen-Berg and T. E. J. Behrens, Diffusion MRI: from quantitative measurement to in-vivo neuroanatomy, 2nd ed. Amsterdam: Elsevier/Academic Press, 2014.

[3] B. T. Thomas Yeo et al., “The organization of the human cerebral cortex estimated by intrinsic functional connectivity,” J. Neurophysiol., vol. 106, no. 3, pp. 1125–1165, Sept. 2011, doi: 10.1152/jn.00338.2011.

[4] J. S. Damoiseaux et al., “Consistent resting-state networks across healthy subjects,” Proc. Natl. Acad. Sci., vol. 103, no. 37, pp. 13848–13853, Sept. 2006, doi: 10.1073/pnas.0601417103.

[5] G. Deco and V. K. Jirsa, “Ongoing Cortical Activity at Rest: Criticality, Multistability, and Ghost Attractors,” J. Neurosci., vol. 32, no. 10, pp. 3366–3375, Mar. 2012, doi: 10.1523/JNEUROSCI.2523-11.2012.

[6] M. W. Cole, T. Ito, D. S. Bassett, and D. H. Schultz, “Activity flow over resting-state networks shapes cognitive task activations,” Nat. Neurosci., vol. 19, no. 12, pp. 1718–1726, Dec. 2016, doi: 10.1038/nn.4406.

[7] C. Seguin, M. P. Van Den Heuvel, and A. Zalesky, “Navigation of brain networks,” Proc. Natl. Acad. Sci., vol. 115, no. 24, pp. 6297–6302, June 2018, doi: 10.1073/pnas.1801351115.

[8] G. L. Baum et al., “Development of structure–function coupling in human brain networks during youth,” Proc. Natl. Acad. Sci., vol. 117, no. 1, pp. 771–778, Jan. 2020, doi: 10.1073/pnas.1912034117.

[9] J. D. Medaglia et al., “Functional alignment with anatomical networks is associated with cognitive flexibility,” Nat. Hum. Behav., vol. 2, no. 2, pp. 156–164, Dec. 2017, doi: 10.1038/s41562-017-0260-9.

[10] X. Jiang et al., “Connectome analysis of functional and structural hemispheric brain networks in major depressive disorder,” Transl. Psychiatry, vol. 9, no. 1, p. 136, Apr. 2019, doi: 10.1038/s41398-019-0467-9.

[11] H. Jiang et al., “Structural–functional decoupling predicts suicide attempts in bipolar disorder patients with a current major depressive episode,” Neuropsychopharmacology, vol. 45, no. 10, pp. 1735–1742, Sept. 2020, doi: 10.1038/s41386-020-0753-5.

[12] S. M. Soman, N. Vijayakumar, P. Thomson, G. Ball, C. Hyde, and T. J. Silk, “Cortical structural and functional coupling during development and implications for attention deficit hyperactivity disorder,” Transl. Psychiatry, vol. 13, no. 1, p. 252, July 2023, doi: 10.1038/s41398-023-02546-8.

[13] A. Zarkali, P. McColgan, L.-A. Leyland, A. J. Lees, G. Rees, and R. S. Weil, “Organisational and neuromodulatory underpinnings of structural-functional connectivity decoupling in patients with Parkinson’s disease,” Commun. Biol., vol. 4, no. 1, p. 86, Jan. 2021, doi: 10.1038/s42003-020-01622-9.

[14] C. J. Honey et al., “Predicting human resting-state functional connectivity from structural connectivity,” Proc. Natl. Acad. Sci., vol. 106, no. 6, pp. 2035–2040, Feb. 2009, doi: 10.1073/pnas.0811168106.

[15] G. Deco, A. Ponce-Alvarez, D. Mantini, G. L. Romani, P. Hagmann, and M. Corbetta, “Resting-State Functional Connectivity Emerges from Structurally and Dynamically Shaped Slow Linear Fluctuations,” J. Neurosci., vol. 33, no. 27, pp. 11239–11252, July 2013, doi: 10.1523/JNEUROSCI.1091-13.2013.

[16] J. Goñi et al., “Resting-brain functional connectivity predicted by analytic measures of network communication,” Proc. Natl. Acad. Sci., vol. 111, no. 2, pp. 833–838, Jan. 2014, doi: 10.1073/pnas.1315529111.

[17] S. F. Kelemen, J. Gõni, S. Pequito, and A. Ashourvan, “A Linear Generative Framework for Structure-Function Coupling in the Human Brain,” July 08, 2025, arXiv: arXiv:2507.06136. doi: 10.48550/arXiv.2507.06136.

[18] J. A. Cummings, B. Sipes, D. H. Mathalon, and A. Raj, “Predicting Functional Connectivity From Observed and Latent Structural Connectivity via Eigenvalue Mapping,” Front. Neurosci., vol. 16, p. 810111, Mar. 2022, doi: 10.3389/fnins.2022.810111.

[19] T. Sarwar, Y. Tian, B. T. T. Yeo, K. Ramamohanarao, and A. Zalesky, “Structure-function coupling in the human connectome: A machine learning approach,” NeuroImage, vol. 226, p. 117609, Feb. 2021, doi: 10.1016/j.neuroimage.2020.117609.

[20] A. Zalesky, T. Sarwar, Y. Tian, Y. Liu, B. T. T. Yeo, and K. Ramamohanarao, “Predicting an individual’s functional connectivity from their structural connectome: Evaluation of evidence, recommendations, and future prospects,” Netw. Neurosci., vol. 8, no. 4, pp. 1291–1309, Dec. 2024, doi: 10.1162/netn_a_00400.

[21] P. Chen et al., “Group-common and individual-specific effects of structure-function coupling in human brain networks with graph neural networks”.

[22] Y. Hong et al., “Structural and functional connectome relationships in early childhood,” Dev. Cogn. Neurosci., vol. 64, p. 101314, Dec. 2023, doi: 10.1016/j.dcn.2023.101314.

[23] H. Yuan, H. Yu, S. Gui, and S. Ji, “Explainability in Graph Neural Networks: A Taxonomic Survey,” July 01, 2022, arXiv: arXiv:2012.15445. doi: 10.48550/arXiv.2012.15445.

[24] R. Ying, D. Bourgeois, J. You, M. Zitnik, and J. Leskovec, “GNNExplainer: Generating Explanations for Graph Neural Networks,” Nov. 13, 2019, arXiv: arXiv:1903.03894. doi: 10.48550/arXiv.1903.03894.

[25] D. Luo et al., “Parameterized Explainer for Graph Neural Network,” Nov. 09, 2020, arXiv: arXiv:2011.04573. doi: 10.48550/arXiv.2011.04573.

[26] H. Yuan, H. Yu, J. Wang, K. Li, and S. Ji, “On Explainability of Graph Neural Networks via Subgraph Explorations,” May 31, 2021, arXiv: arXiv:2102.05152. doi: 10.48550/arXiv.2102.05152.

[27] S. Lundberg and S.-I. Lee, “A Unified Approach to Interpreting Model Predictions,” Nov. 25, 2017, arXiv: arXiv:1705.07874. doi: 10.48550/arXiv.1705.07874.

[28] J. Royer et al., “An Open MRI Dataset For Multiscale Neuroscience,” Sci. Data, vol. 9, no. 1, p. 569, Sept. 2022, doi: 10.1038/s41597-022-01682-y.

[29] R. R. Cruces et al., “Micapipe: A pipeline for multimodal neuroimaging and connectome analysis,” NeuroImage, vol. 263, p. 119612, Nov. 2022, doi: 10.1016/j.neuroimage.2022.119612.

[30] T. N. Kipf and M. Welling, “Semi-Supervised Classification with Graph Convolutional Networks,” Feb. 22, 2017, arXiv: arXiv:1609.02907. doi: 10.48550/arXiv.1609.02907.

[31] S. S. Haykin, Neural networks: a comprehensive foundation, Nachdr. New York, NY: Macmillan [u.a.], 1995.

[32] M. Sundararajan, A. Taly, and Q. Yan, “Axiomatic Attribution for Deep Networks,” June 13, 2017, arXiv: arXiv:1703.01365. doi: 10.48550/arXiv.1703.01365.

[33] C. Seguin, Y. Tian, and A. Zalesky, “Network communication models improve the behavioral and functional predictive utility of the human structural connectome,” Netw. Neurosci., vol. 4, no. 4, pp. 980–1006, Jan. 2020, doi: 10.1162/netn_a_00161.

[34] C. Seguin, S. Mansour L, O. Sporns, A. Zalesky, and F. Calamante, “Network communication models narrow the gap between the modular organization of structural and functional brain networks,” NeuroImage, vol. 257, p. 119323, Aug. 2022, doi: 10.1016/j.neuroimage.2022.119323.

[35] D. J. Lurie et al., “Questions and controversies in the study of time-varying functional connectivity in resting fMRI,” Netw. Neurosci., vol. 4, no. 1, pp. 30–69, Jan. 2020, doi: 10.1162/netn_a_00116.

[36] A. Eichenbaum, I. Pappas, D. Lurie, J. R. Cohen, and M. D’Esposito, “Differential contributions of static and time-varying functional connectivity to human behavior,” Netw. Neurosci., vol. 5, no. 1, pp. 145–165, Jan. 2021, doi: 10.1162/netn_a_00172.

[37] P. B. Kemmer, Y. Wang, F. D. Bowman, H. Mayberg, and Y. Guo, “Evaluating the Strength of Structural Connectivity Underlying Brain Functional Networks,” Brain Connect., vol. 8, no. 10, pp. 579–594, Dec. 2018, doi: 10.1089/brain.2018.0615.

[38] C. Zhou, L. Zemanová, G. Zamora, C. C. Hilgetag, and J. Kurths, “Hierarchical Organization Unveiled by Functional Connectivity in Complex Brain Networks,” Phys. Rev. Lett., vol. 97, no. 23, p. 238103, Dec. 2006, doi: 10.1103/PhysRevLett.97.238103.

[39] B. Corominas-Murtra, J. Goñi, R. V. Solé, and C. Rodríguez-Caso, “On the Origins of Hierarchy in Complex Networks,” Proc. Natl. Acad. Sci., vol. 110, no. 33, pp. 13316–13321, Aug. 2013, doi: 10.1073/pnas.1300832110.

[40] M. R. Arbabshirani et al., “Impact of autocorrelation on functional connectivity,” NeuroImage, vol. 102, pp. 294–308, Nov. 2014, doi: 10.1016/j.neuroimage.2014.07.045.

[41] R. F. Betzel et al., “Generative models of the human connectome,” NeuroImage, vol. 124, pp. 1054–1064, Jan. 2016, doi: 10.1016/j.neuroimage.2015.09.041.

[42] R. D. Markello and B. Misic, “Comparing spatial null models for brain maps,” NeuroImage, vol. 236, p. 118052, Aug. 2021, doi: 10.1016/j.neuroimage.2021.118052.

[43] E. S. Finn et al., “Functional connectome fingerprinting: identifying individuals using patterns of brain connectivity,” Nat. Neurosci., vol. 18, no. 11, pp. 1664–1671, Nov. 2015, doi: 10.1038/nn.4135.

[44] B. Chiêm, K. Abbas, E. Amico, D. A. Duong-Tran, F. Crevecoeur, and J. Goñi, “Improving Functional Connectome Fingerprinting with Degree-Normalization,” Brain Connect., p. brain.2020.0968, Aug. 2021, doi: 10.1089/brain.2020.0968.

[45] D. A. Fair et al., “Functional Brain Networks Develop from a ‘Local to Distributed’ Organization,” PLoS Comput. Biol., vol. 5, no. 5, p. e1000381, May 2009, doi: 10.1371/journal.pcbi.1000381.

[46] A. Fornito, A. Zalesky, and M. Breakspear, “The connectomics of brain disorders,” Nat. Rev. Neurosci., vol. 16, no. 3, pp. 159–172, Mar. 2015, doi: 10.1038/nrn3901.

[47] M. P. Van Den Heuvel et al., “Abnormal Rich Club Organization and Functional Brain Dynamics in Schizophrenia,” JAMA Psychiatry, vol. 70, no. 8, p. 783, Aug. 2013, doi: 10.1001/jamapsychiatry.2013.1328.

[48] Y. Stern, “Cognitive reserve in ageing and Alzheimer’s disease,” Lancet Neurol., vol. 11, no. 11, pp. 1006–1012, Nov. 2012, doi: 10.1016/S1474-4422(12)70191-6.

[49] F. Agosta, M. Pievani, C. Geroldi, M. Copetti, G. B. Frisoni, and M. Filippi, “Resting state fMRI in Alzheimer’s disease: beyond the default mode network,” Neurobiol. Aging, vol. 33, no. 8, pp. 1564–1578, Aug. 2012, doi: 10.1016/j.neurobiolaging.2011.06.007.

